# The *PIN2* ortholog in barley modifies root gravitropism and architecture without impacting the shoot

**DOI:** 10.1101/2024.03.28.587117

**Authors:** Zachary Aldiss, Hannah Robinson, Yasmine Lam, Richard Dixon, Peter Crisp, Ian Godwin, Andrew Borrell, Lee Hickey, Karen Massel

**Affiliations:** Queensland Alliance for Agriculture and Food Innovation, The University of Queensland, Brisbane, QLD, Australia; InterGrain Pty Ltd, Perth, WA, Australia; School of Agriculture and Food Sustainability, The University of Queensland, Brisbane, QLD, Australia; Queensland Alliance for Agriculture and Food Innovation, The University of Queensland, Warwick, QLD, Australia

**Keywords:** Biotechnology, Auxin, CRISPR, Genome Engineering, Allometry

## Abstract

Roots provide the critical interface where plants acquire nutrients and water, but our limited understanding of the genetic controls modulating root system architecture (RSA) in crop species constrains opportunities to develop future cultivars with improved root systems. However, there is vast knowledge of root developmental genes in model plant species, which has the potential to accelerate progress in crops with more complex genomes, particularly given that genome editing protocols are now available for most species. *PIN-FORMED2* (*PIN2*) encodes a root specific polar auxin transporter, where its absence resulted in roots being unable to orient themselves using gravity, producing a significantly wider root system. To explore the role of *PIN2* in a cereal crop, we used CRISPR/Cas9 editing to knockout of *PIN2* in barley (*Hordeum vulgare*). Like Arabidopsis, the roots of barley *pin2* loss-of-function mutants displayed an agravitropic response at seedling growth stages, resulting in a significantly shallower and wider root system at later growth stages. Notably, despite the significant change in RSA, there was no change in shoot architecture or total shoot biomass. We discuss the future challenges and opportunities to harness the *PIN2* pathway to optimise RSA in crops for a range of production scenarios without a shoot trade-off.

## INTRODUCTION

The uptake of water and nutrients from the soil is critical to support crop growth and productivity. With a rapidly growing global population and the increasing effects of climate change, there is a greater demand for crops that can produce more grain with less resources. Breeding crops with enhanced sustainability relies on access to genetic variation for morphological and physiological traits that influence resource-use efficiency and adaptation to abiotic stress (Vanhala et al. 2004; Cronin et al. 2007). While there is still limited understanding of the genetics modulating crop development, significant exploration has been performed in model species Arabidopsis, which provides a powerful resource bank of candidate genes to surgically explore the various mechanisms of crop physiology and development. So far, the crop research community has made good progress to investigate the translation of gene targets identified in model species for above-ground architecture (Ahmar et al. 2024; Huang et al. 2021), leaving many promising targets for root system architecture (RSA) as a novel opportunity to improve nutrient- and water-use efficiency. RSA has important functional implications as it is the primary interface for soil nutrient and water acquisition, along with the physical anchoring of the plant and providing resilience to lodging (Bishopp and Lynch 2015). Among the most important determinants of RSA is the root growth angle (RGA), which is the angle at which the roots grow into the soil.

A key mechanism influencing RGA is how the roots respond to gravity, where a stronger gravitropic response elicits a steeper rooting angle and is often associated with improved drought adaptation through improved access to deep-soil water (Kitomi et al. 2020; Ho et al. 2004). Root gravitropism is regulated by the phytohormone auxin (indole-3-acetic acid or IAA), a ubiquitous and important chemical responsible for translating external changes to the environment into a physiological response by the plant, allowing for rapid adaptation to its environment (Blilou et al. 2005). This process is maintained by polar auxin transporters, establishing asymmetrical organ-specific and organism-wide concentration gradients that regulate cell elongation to enable root growth along the gravity vector. Removing the root cap through genetic ablation results in loss of gravity sensing in the roots and increased lateral rooting, a response consistent with auxin insensitivity or inhibition (Tsugeki and Fedoroff 1999; Mirza et al. 1984). The starch-statolith hypothesis is the prevailing theory on how the root cells sense gravity, where dense starch granules named amyloplasts sediment on one side of the cell aligned with the gravity vector, triggering a signalling cascade (Stanga et al. 2011). While still poorly understood, it is thought that this cascade involves the rearrangement of auxin transporters to reestablish the auxin maximum at the root tip.

Auxin distribution is facilitated by three major classes of transporters: the PIN-FORMED (PIN) efflux carrier proteins, the ATP-binding cassette (ABC)-B/multi-drug resistance/P-glycoprotein (ABCB/MDR/PGP) subfamily of ABC transporters, and the AUXIN1/LIKE-AUX1 (AUX/LAX) influx carrier proteins (Yue et al. 2015). The ABCD/MDR/PGP transporters are a large family of transporters with diverse roles in the transport of a variety of substrates, where some studies have demonstrated their involvement in both influx and efflux with auxin (Geisler and Murphy 2006). Members of the AUX/LAX family facilitate auxin entry into cells through passive diffusion, leading to unidirectional distribution of auxin due to their asymmetric positioning on the plasma membrane (PM) (Swarup et al. 2001). The most well described polar auxin transporters are the PIN proteins responsible for auxin efflux.

There are two main types of *PIN* genes: the ‘short’ PINs and ‘long’ PINs based on the size of the intracellular hydrophilic loop domain that separates the flanking transmembrane domains (Luschnig and Vert 2014). Short PINs play an essential role in maintaining intracellular auxin concentrations by regulating transport between the endoplasmic reticulum and cytosol (Geisler and Murphy 2006), whereas long PINs regulate the efflux of auxin out of the cell through asymmetrical localization on the PM. This localisation changes in response to environmental stimuli by cycling the receptors between the PM and the endoplasmic reticulum (Kirschner et al. 2018). This is established through unique and regulated polar localization of these transporters along the plasma membrane, allowing directional transport of auxin and forming unique local distributions for organ specific, and organism wide responses (Blilou et al. 2005). Arabidopsis has a total of nine known PIN transporters, whereas higher order species such as cereals often have an increased number of PIN transporters. For example, barley has 13 putative PIN transporters with Hv-PIN1 and Hv-PIN2 being homologous to Arabidopsis long PINs At-PIN1 and At-PIN2. In contrast, Hv-PIN5, Hv-PIN8 and Hv-PIN9 exhibit short hydrophilic regions. While monocots do not encode At-PIN3, At-PIN4 and At-PIN7 homologues, barley uniquely has four Hv-PIN10s which display atypical structure of the hydrophilic loop being non-central (Hv-PIN10b2) or missing entirely (Hv-PIN10a, Hv-PIN10b1 and Hv-PIN10b3) (Kirschner et al. 2018).

The developmental plasticity of plants is underpinned by direct and indirect effects. Direct effects controlling organ formation are primarily the consequence of local hormone interactions. Indirect effects can be seen in the dynamic balance of allometry where growth of one organ influences another. For example, the canopy has an allometric relationship with the root system, where increases in the canopy biomass correlates with increased root biomass (Borrell et al. 2022). As *PIN2* expression is almost exclusively within the roots, the possibility of root specific architectural changes, independent of above- and below-ground allometry, provides an interesting subject for investigation (Müller et al. 1998). PIN2 is a known regulator of root gravitropism; it localizes to the apical side of epidermal cells and mediates the auxin flow along the lower side of the root from the tip toward the elongation zone where the auxin response is activated and inhibits cell elongation (Casimiro et al. 2001). In *Arabidopsis, pin2* mutants show defective root gravitropism because of the mutants’ inability to accumulate auxin in the roots, a loss of curving in RSA formation.

While the PIN2 pathway is well-studied in model species, knowledge of the extent to which *PIN2* function is preserved across species, particularly monocots, is lacking. In this study, we explore the function and potential of targeted manipulation of the *PIN2* orthologue (Hv*-PIN2*) in the major cereal crop, barley (*Hordeum vulgare*). We created a knockout in barley using CRISPR/Cas9 genome editing and adopted a wholistic plant phenotyping approach that measured a range of root and shoot traits across growth stages to provide new insights of the function and potential utility of genome editing to support crop improvement.

## MATERIALS AND METHODS

### CRISPR/Cas9 editing of *PIN2* in barley

Genome editing of *PIN2* (*HORVU7Hr1G110470*) in barley was performed using *cv.* Golden Promise via particle bombardment following the tissue culture protocol described in Harwood et al. (2009), with modifications and gene editing strategy explained in Massel et al. (2022). Wild type (WT) embryos that did not go through transformation remained on non-selective regeneration media. Regenerants that produced shoots at least 2cm long were subsequently transferred to rooting media for planting.

Methods for selection of gRNA are described in Massel et al. (2022). Using those parameters, two gRNAs (G1: 5’-GTGGCCAACAAGTTCAAGGG-3’, G2: 5’-GCCGGTAGTCCATGGCGTAG-3’) targeting coding regions of the *PIN2* orthologue were selected and cloned into the plasmid system. Following isolation of barley immature embryos, transformation was performed via microprojectile co-bombardment using biolistic techniques (Liu and Godwin 2012) using the particle inflow gun delivery system with modifications to optimize the system for barley (Liu et al. 2014). Immature embryos were bombarded approximately 5-7 days post isolation with approximately 25-30 embryos per shot. During bombardment, the vacuum pressure remained at −90kHa with helium being shot at a pressure of 800kPa.

Following successful tissue culture, leaf tissue samples were collected from the putatively edited lines for DNA extraction, PCR and DNA Sanger sequencing to identify the nature of the edit. PCR was performed using MyTaq (Bioline) DNA Polymerase in reactions containing 30ng of DNA template, 1X MyTaq Buffer, 0.5uM of the forward and reverse primers (Table 1) and run following the manufacturer’s instructions. PCR reactions were run on 1.2% TAE agarose gel with a standard 1kb ladder (New England BioLabs) to confirm formation of products. PCR clean-up was performed with ExoSAP-IT™ (Thermo Fisher Scientific) before being sent for Sanger sequencing. Genotypic analysis is performed using CRISPR chromatogram analysis tool Synthego (Synthego ICE Analysis tool v3), along with sequence alignment and chromatograph analysis. Transmembrane prediction software TMHMM was used to estimate resultant impact on protein function (Krogh et al. 2001).

**Table 1.**
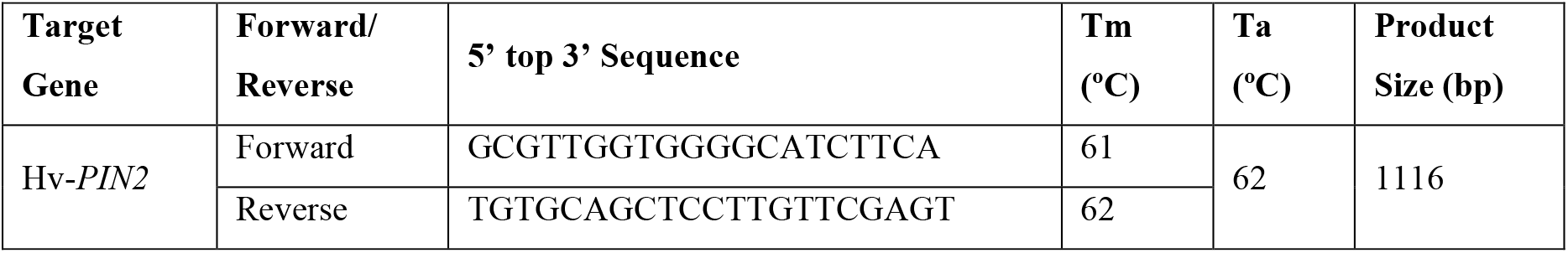
Primers used in PCR to detect presence of gene and potential edits following transformation.

### Evaluating gravitropic response

To visualise and monitor the gravitropic response, seeds were germinated in glass chambers (210 x 297 mm) lined with black serviettes that were watered and rolled flat to remove air bubbles prior to placing seeds. The developing plants were grown in darkness at 25 ℃. After 4 days, images of the roots were captured and then chambers were rotated by 90°. Root images were captured at 0, 4, 6, 8 and 24 hours after rotation, with the root position marked at each time point. The responsiveness of the root system to gravity was quantified by measuring the angle of the root tips for seminal roots and the radicle over time (Kirschner et al. 2021).

For data analysis, the root angle values for all root tips were measured in relation to the horizontal, with the angle following rotation set to 0°. All root angle values were obtained with the software ImageJ (Schneider et al. 2012). The average root angle at each time point and change in root angle over time were calculated and compared using a two-tailed t-test.

### Phenotyping seminal root angle

The ‘clear pot’ method, routinely applied in wheat and barley root research (Richard et al. 2015; Robinson et al. 2016), was used to evaluate the *pin2* knockout and WT barley lines for seminal root angle and number at the seedling stage. Seminal root angle is considered a useful proxy trait that can be assayed in a high-throughput manner and is often associated with the root angle of mature plants in the field (Alahmad et al. 2019; Voss-Fels et al. 2018). Plants were watered daily and housed in a temperature controlled PC2 glasshouse, with 22℃ /17℃ (day/night) temperature cycle and 18hr/6hr (day/night) photoperiod.

All edited lines were assessed and compared to the WT as well as a tissue culture control plant (i.e. unedited line regenerated from tissue culture). The edited lines screened included two mutants (*pin2-1* and *pin2-2,* Figure 1). Seeds were sown in 4 L ANOVA pots of 200 mm diameter and 190 mm in height using a randomised design ensuring 12 reps for each line, with a maximum of 24 seeds per pot. Seeds were sown at a depth of 2 cm at a 45° angle, with the embryonic axis of each seed oriented downwards, every 2.5 cm along the transparent wall of the pot to result in 24 barley seeds per pot (Robinson et al. 2016). UQ-23 potting mix was used, containing 70% composted pine bark 0–5 mm, 30% cocoa peat.

**Figure 1.**
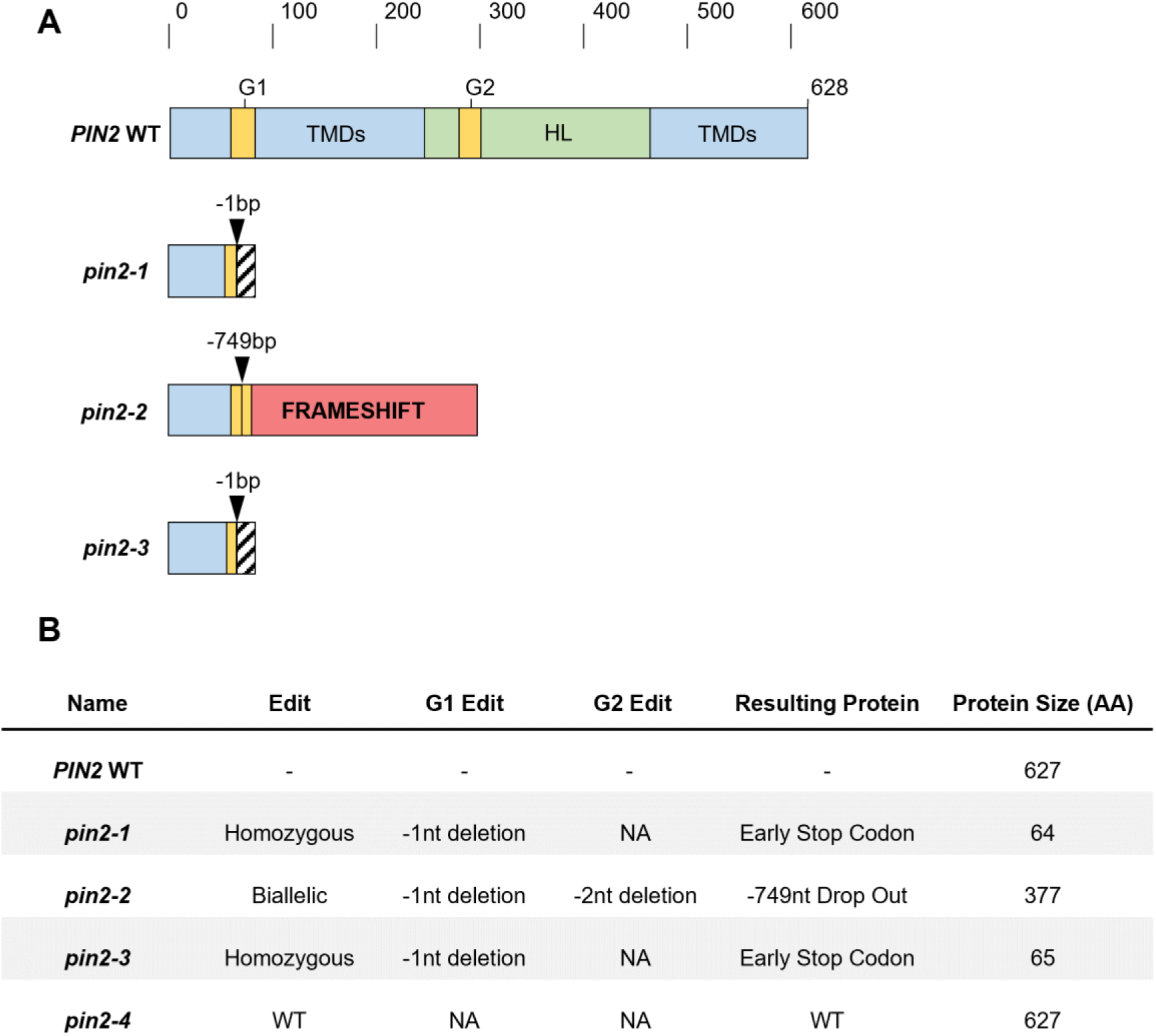
Summary of barley *pin2* knock-out lines created using genome editing and resultant protein structure changes. **A)** Map of alleles for the long-looped *PIN* gene showing generalised edits to protein sequence. Scale represents the number of amino acids. Specifically, *pin2-1* displayed an early stop codon following G1, *pin2-2* displayed a 749bp drop out along with a frame shift for the remaining coding region, and *pin2-3* showed the same editing event −1nt deletion at G1 and was homozygous. Notably, *pin2-4* displayed no genome editing at *PIN2* and was treated as the tissue culture (TC) control for future experiments. **B)** Outcomes table of transgenics, with the number of alleles edited at each guide location and the resulting PIN2 protein. G1 and G2 = Guide sites one and two for CRISPR/Cas9 genome editing, TMD = Transmembrane Domain, HL = Hydrophilic Loop.

The pots were thoroughly watered the day before sowing and no additional water was provided, which ensured the orientation of seeds did not shift during germination. After sowing, each clear pot was placed inside a black pot of the same dimension to prevent light from influencing root development. Images of roots were captured at both three and five days after sowing using a camera (iPhone 13 Pro Max). Analysis for seminal root angle and number was determined using ImageJ (Schneider et al. 2012), measuring the average deviation angle of the first pair of seminal roots to the vertical plane (Robinson et al. 2016; Richard et al. 2015; Christopher et al. 2013).

### Evaluating root system architecture in rhizoboxes

The RSA of the *pin2* and WT lines were assessed in a series of rhizobox experiments. RSA was assessed by determining the root angle, total biomass, and root density at various depths. This set-up uses wooden boxes lined with a plastic sheet and placed upright on a bench with capillary matting at the base of the bench. The first two experiments were conducted using the rhizobox platform described by Kang et al. (2024). The chambers were 30 cm (width) x 5 cm (thickness) x 80 cm (depth) and were filled with potting mix containing 6 g/L of Osmocote mixed in thoroughly. Plants were watered daily and housed in a temperature controlled PC2 glasshouse, with 22℃/17℃ (day/night) temperature cycle and standard long-day conditions using a 18hr/6hr (light/dark) photoperiod. Two different planting densities were tested, first with three plants per chamber, and second with one plant per chamber. In the first experiment, six to nine seeds were evenly dispersed throughout the width of the chamber near the surface of the potting mix (5 cm depth). After emergence each chamber was thinned down to three plants per box. In the second rhizobox experiment, a total of three seeds were sown (depth of 5 cm) per rhizobox with one plant retained after emergence. The experiments in these rhizoboxes were harvested after four weeks, as the root systems had almost reached the base of the chambers. Two days prior to harvesting the plants were heavily watered and were not watered the day prior to harvest. Upon harvesting, in both rhizobox experiments, each chamber was divided into three equal segments (26 cm depth) to represent ‘upper’, ‘middle’ and ‘deep’ layers. The potting media-containing roots sampled from each section was collected and bagged individually. Each root segment was hand washed to remove potting mix, placed in a paper bag and dried at 80°C before weighing.

While the plants grew well in the 30 cm width boxes, the narrow width could not accommodate the outgrowth of the mutant root system as the roots reached the sides of the chamber early in the plant growth cycle resulting in either ‘training’ of the roots downwards or ‘bouncing’ off the wall and back into the substrate. This reduced the ability to accurately quantify differences in RSA. To accommodate the wider root system, ‘double width’ rhizoboxes (60 cm x 5 cm x 80 cm) were constructed to improve the visualisation of the entire RSA.

In the ‘double width’ rhizboxes, plants were phenotyped after 42 days of growth, which enabled the analysis of root systems that were unimpeded and more extensively developed. The same potting mix, growing and plant maintenance parameters were used. For the double width rhizobox experiment, a total of three seeds were sown (depth of 5 cm), which were later thinned to one plant per rhizobox. Thus, it was anticipated that growing barley plants in these wider rhizoboxes would provide a clearer visualisation of the root system branching and distribution down to a depth of 80 cm and root:shoot biomass partitioning in a setting that more closely resembled field conditions. The double width rhizobox was analysed by separating the chamber and image into nine biomass sections (3 x 3 sections of 20 cm x 26 cm). As the root structures of the individual plants in the wide rhizoboxes could not be separated, the root biomass measurements represented the total root biomass produced by the three plants.

Experiments in both rhizobox platforms were laid out as randomised complete block designs for the WT and *pin2* edited line (*pin2-1*) using six to eight reps per genotype. To synchronise germination, a cold treatment was performed by placing the seed in a petri dish lined with a moist paper towel overnight at 4℃, then plates were moved to room temperature (22℃) for 24hrs prior to sowing. The rhizoboxes were well-watered daily by manually adding water to the top until drainage was observed from the bottom. During the experiment, a leaf sample from each plant in the box was taken for DNA extraction to confirm the genotype of the plants.

The root system was photographed for future analysis via RhizoVision Explorer to quantify proportion of roots with shallow, medium and steep root angles grouping by those less than 30, less than 60 and less than 90 degrees from the medial axis (Seethepalli et al. 2020). To investigate potential changes in above-ground biomass the entire shoot sample for each plant was removed and placed in individually labelled bags.

### Evaluating above-ground traits at physiological maturity

To further explore any shoot trade-offs associated with *pin2* in barley, the *pin2* edited lines (*pin2-1, pin2-2*) and WT were grown under controlled conditions and evaluated at physiological maturity. Plants were sown in 1 L pots at a density of one plant per pot using eight reps per genotype arranged in a randomised complete block design. Plants were grown under stress-free conditions using a temperature regime of 22℃/17℃ (day/night) and a standard long-day photoperiod of 18hr/6hr (light/dark). At maturity, each plant was partitioned into spikes, stems with leaves and grains. Plant height, spike length and the number of seed per spike were recorded for the primary tiller of each plant. The total number of fertile tillers (grain producing tillers) and days to flowering were recorded for each plant.

### Analysis of root and shoot phenotype data

To analyse data from the gravitropism experiment, the root angles obtained by *pin2* edited line and WT line were compared using parametric t-tests and one-way ANOVAs with multiple comparisons where H_0_ > 0.05 using GraphPad Prism (v9.3.1).

All root and shoot data collected in glasshouse phenotyping experiments, including clear pot, rhizobox and maturity studies, were analysed using a linear mixed model (LMM) to account for spatial variation and maximise the genetic variance captured in each trait specific model. The LMM was fitted with the following equation:

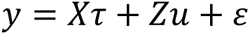

where X and Z are design matrices associated with fixed (*τ*) and random (*u*) effects and *ε* is the residual errors. A custom linear mixed model was fit to each trait measured across the glasshouse root and shoot experiments, with the best model fit determined by the Wald test statistics (for fixed effect terms only), the highest REML loglikelihood value and the smallest Akaike Information Criterion (AIC) (Kenward and Roger 1997; Akaike 1974). The specific design terms fitted as random and fixed effects in the LMM for each trait are detailed in Table S1. The auto-regressive residual variance structure in the order of 1 was found to fit best across all traits and was thus fitted across all trait LMMs. A generalised measure of broad sense heritability was calculated for each trait to quantify the proportion of phenotypic variance that is attributed to genetic effects. This was determined using the following equation:

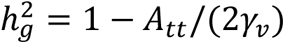

where *A*_*tt*_ is the mean predicted error variance and *γ*_**ν**_ is the genetic variance (Cullis et al. 2006). The LMMs were fit using ASReml-R in the R software environment (R Development Core Team, 2023), and Best linear unbiased predictions (BLUPs) calculated from the variance components of the model (Butler et al. 2017). Figures displaying and comparing trait BLUPs between *pin2* edited lines and the WT were created using the ggplot package in R (Wickham et al. 2016).

## RESULTS

### Gene edited *pin2* knockout lines in barley

The *PIN2* orthologue in barley was identified using Phytozome, showing 76.8% protein homology and has the highest similarity to *Sorghum bicolor* (Sb-*PIN11*) and *Oryza Sativa* (Os-*PIN2*) orthologues of At-*PIN2* (Goodstein et al. 2012, Table S2). Four stable transgenics were created that contained both plasmids. Of these, two lines were shown to contain edits in at least one of the target sites, and will be described as *pin2-1* and *pin2-2*. Sanger sequencing was used to verify the editing outcomes, where *pin2-1* has a homozygous −1nt deletion, resulting in a frame-shift of the coding region. This leads to a truncated peptide fragment that only contains 64/627 of the WT protein sequence before stopping. Similarly, *pin2-2* was compound heterozygous, edited at both sites, where one allele was a −749bp deletion in the sequence between the two guide sites resulting in a frameshift mutation with 65/627 of the original peptides and 377 amino acids before stopping, while the other allele only had the G1 edit causing an early stop codon. Segregation for the homozygous mutant resulted in *pin2-3,* an edited line with the G1 edit of *pin2-2* resulting in a biallelic early stop codon producing a peptide 65/627 of the original sequence. One line showed no evidence of editing at target sites and was described as *pin2-4.* Editing efficiency was approximately 75% at G1 and 25% at G2. Transmembrane prediction using TMHMM displayed a loss of the transmembrane domains following the edit along with the central hydrophilic loop, domains that are essential to the function of PIN proteins (Figure S1). A summary of transgenics can be seen in Figure 1.

### *PIN2* controls root gravitropism in barley

The rate of change in root angle over time is representative of both gravitropic response and strength of its response. The roots of *pin2-2* (n=7) lacked a gravitropic response, where the roots continued to grow from their initial sprouting (Figure 2). On the other hand, WT (n=8) roots showed a linear rate of change to root angle towards the gravity vector at an average of 0.83 °/hr. The shoots appeared to respond to gravity, with both WT and *pin2-2* curving upwards following rotation.

**Figure 2.**
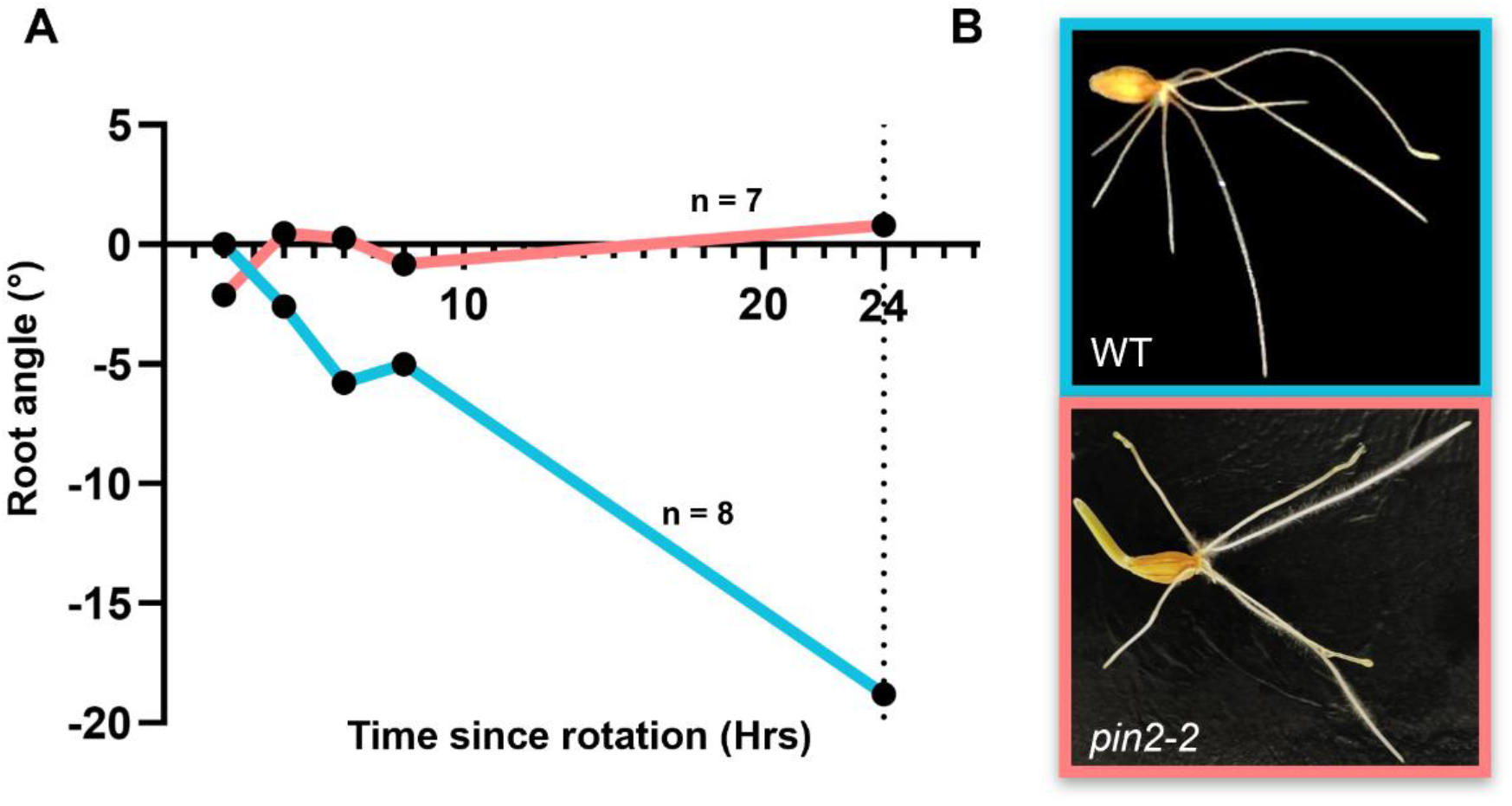
Barley *pin2* lines display root-specific agravitropism at the seedling stage. **A)** Change in root angle over 24 hours from seedling gravitropic assay conducted in glass panels. Root angle was measured from the radicle in relation to the horizontal axis, and measurements were taken every two hours for the first eight hours, then after 24 hours (n=7). The derivative of root angle over time was taken as a standardized measurement for strength of gravitropic response. **B)** Images of WT and *pin2-2* seedlings captured 24 hours after rotation.

**Figure 3.**
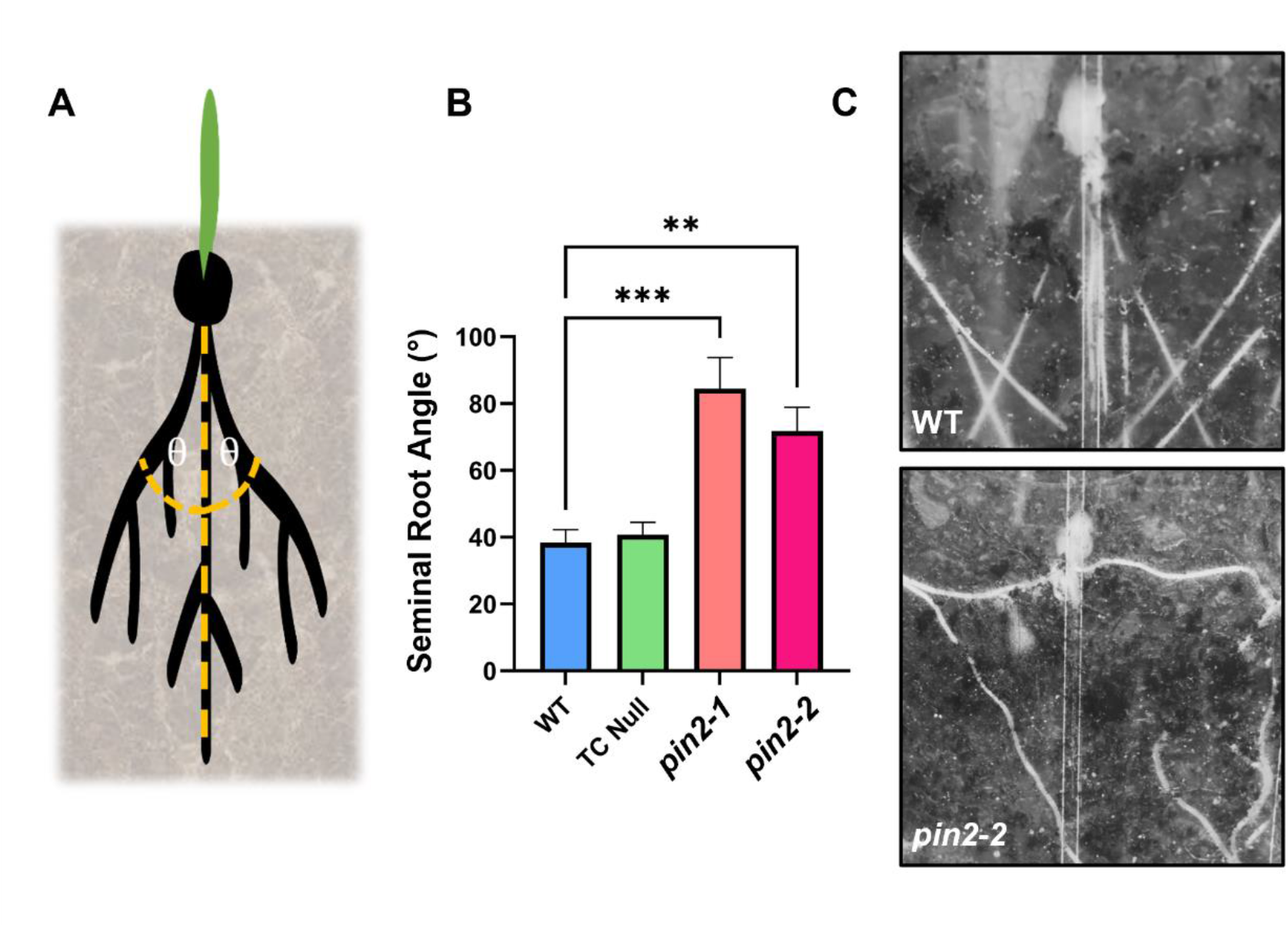
Barley *pin2* knock-out lines display a significantly wider seminal root angle. **A)** Diagram illustrating how the measurements for seminal root angle were recorded in this study. Measurements were recorded in ImageJ for the first pair of seminal roots to the vertical axis and the two angles summed to give a representative measurement of root angle. **B)** Comparison of seminal root angle phenotypes for Golden Promise wild type (WT) carrying *PIN2*, tissue culture (TC) null (*pin2-4*) and transgenic knock-out lines for *pin2* (*pin2-1* and *pin2-2*). **C)** Representative images of seminal root angle displayed by WT and *pin2-2* seedlings using the clear pot method.

### Agravitropic roots produced altered root growth angle

To determine if Hv-*PIN2* plays a role in root growth orientation, the seminal root angle was evaluated using the clear pot system. This was performed on two homozygous knockout lines, *pin2-1* (n=9) and *pin2-2* (n=7), alongside a tissue culture null (n=11) and WT (n=16) as controls. The seedlings were imaged at days 3, 5 and 12. Both *pin2-1* and *pin2-2* displayed a significantly wider seminal root angle, from 38° in WT to 85° and 72° between *pin2-1* and *pin2-2* respectively, displaying similar patterns of agravitropism as those seen in the previous gravitropism assay, where roots appear ‘aimless’, growing in the direction that they emerged (Fig 2B).

### Barley *pin2* knockout lines display wider and shallower root architecture

To explore the role of *PIN2* on RSA, a series of rhizobox experiments were conducted, which compared *pin2-1* and the WT. Initial phenotyping was conducted using narrow rhizoboxes at two planting densities: 1 plant per box (15 cm row spacing) and 3 plants per box (5 cm row spacing) at 28 days after sowing (DAS). In the lower-density experiment, results revealed that *pin2-1* produced approximately 40% more root biomass in the upper section of the rhizoboxes compared to the WT (P<0.05), yet no difference in shoot biomass was detected (P>0.1, Figure S2). In the high-density experiment, *pin2-1* showed a reduction in root biomass of approximately 30% (P=0.07) and no change to above ground biomass (P>0.1). The major difference in root distribution was observed in the lower section (S3), where *pin2-1* produced significantly less root biomass in both low- and high-density experiments (Figure S2). As a result of the reduction in root biomass, *pin2-1* showed a significantly lower root:shoot ratio (RSRatio) compared to the WT at 1- and trending reduction at 3-planting density (P<0.001, P=0.057).

It was evident that the narrow width of the rhizoboxes in the first set of experiments constrained our ability to accurately phenotype RSA of the *pin2* mutants as the roots hit the sides and were trained downward (Figure S2B). Thus, rhizoboxes with double the width were used to subsequently phenotype the lines in a scenario where root growth was less impeded. In the wider rhizoboxes, *pin2-1* exhibited a wider and shallower root system, as demonstrated by the increased root biomass produced in the upper wide sections (S1 and S3; P<0.05, Fig 4A). The upper middle section (S2) (i.e. directly below the base of the plant) showed no significant change in root biomass whereas the middle section (S5) showed a significant reduction in root biomass (P<0.05). In the wide rhizoboxes there was no significant change to above-(P>0.1) or below-ground biomass (P>0.1) in *pin2-1* compared to the WT. However, image analysis of the whole root system using RhizoVision Explorer showed that *pin2-1* displayed a higher proportion of roots classified as ‘shallow’ (P<0.05), indicative of shallow rooting behaviour (Fig 4D).

**Figure 4.**
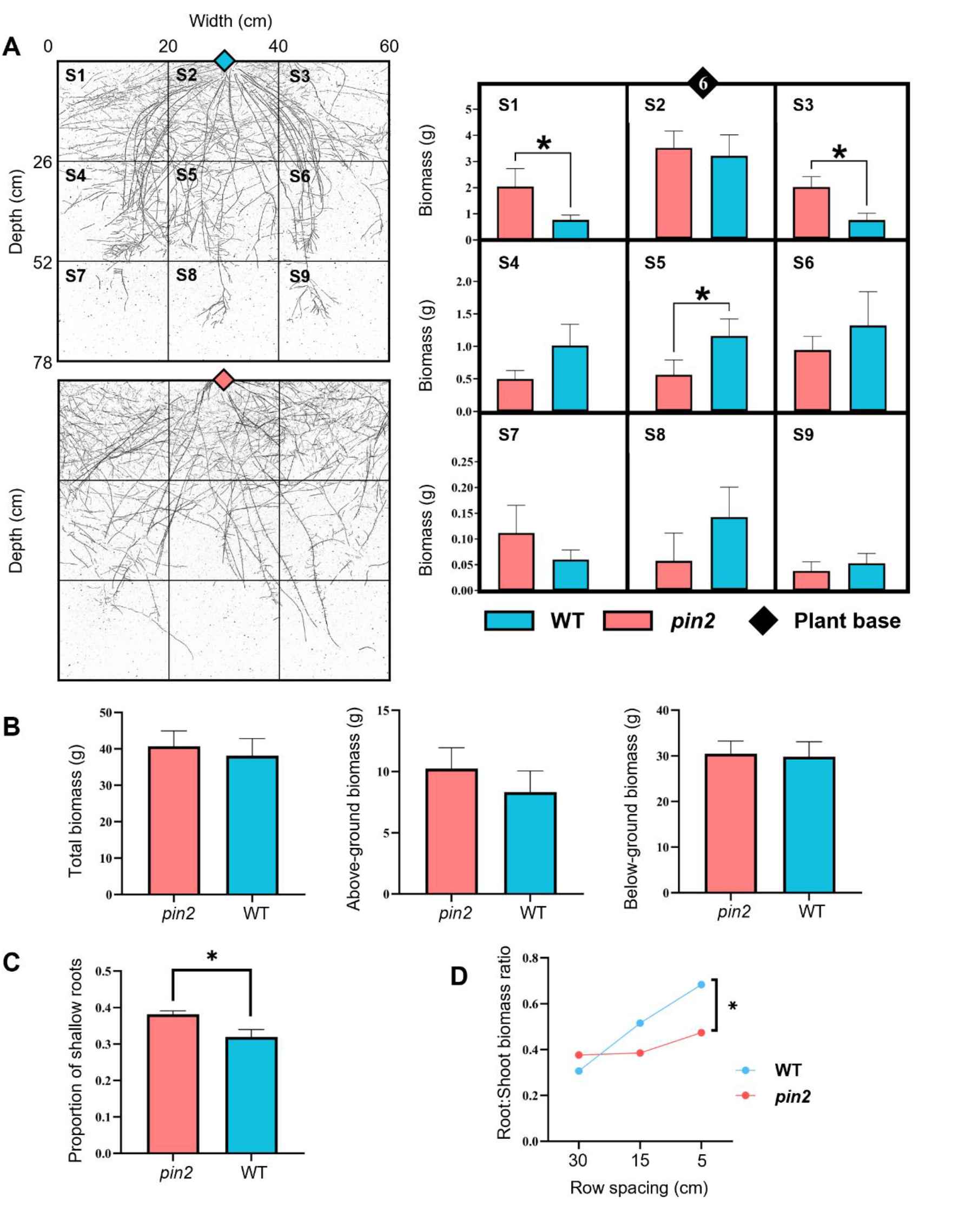
PIN2 in barley regulates downward root growth leading to steeper and narrower root architecture. **A)** Spatial root system configuration of the wild type (WT) and *pin2* barley lines in the wide rhizobox chambers. Left images show representative images of the root systems for both the WT (Blue) and *pin2-1* (Red). Solid lines indicate each section (S1-S9). The dry root biomass (g) produced in each section is displayed as bar charts on the right. **B)** Biomass of total, above-ground and below-ground biomass for both genotypes as an indicator of changes to priority in resource allocation. No significant differences between WT and *pin2* across all sections were observed. **C)** The proportion of total roots classified as ‘shallow’ roots based on image analysis using RhizoVision Explorer, where the orientation of each root in the whole rhizobox surface area is allocated to either shallow (<30°), medium (<60°) or steep angle (<90°). No significant differences were detected for steep and medium angle classes, but there was a significant increase in shallow angle roots displayed by *pin2*. **D)** The root:shoot ratio for WT and *pin2* genotypes grown at three different row spacings indicative of planting density. Error bars represent standard error of the spatially adjusted mean for A, B and C with standard error of mean for D. Significance was calculated by non-parametric t-test. N=6. * = p<0.05.

### *PIN2* has a limited role in above-ground development

To explore any allometric relationships or potential above-ground trade-offs, canopy branching was measured. A seedling assay evaluated knockout lines (*pin2-1* and *pin2-2*) and WT control (n=14), where numerous shoot traits were recorded after 30 days of growth. Leaf areas of the first five leaves showed consistent linear growth over time, with no significant differences between WT and edited lines (P>0.05).

Below-ground dry weight was also calculated after washing the soil off the seedling roots, and demonstrated reduced root mass, indicating a decrease in resource allocation to the roots in mutants with a significant reduction in root:shoot biomass ratio (Fig 4C). Below ground biomass was 58 g for WT with 44 g and 28 g for *pin2-1* and *pin2-2* while above-ground biomass was 77 g for WT and 64 g and 41g for *pin2-1* and *pin2-2* respectively. The root-to-shoot biomass ratio was significantly altered in *pin2* with from 0.6 in WT to 0.45 representing a general reduction in root biomass of 15% (P<0.01, Figure S3).

Across both rhizobox trials, no significant difference in canopy biomass was detected despite the significantly reduced root biomass. Early vigor measurements conducted on the three plant per rhizobox trial revealed no significant difference in tiller number (P>0.1) or leaves per tiller (P>0.1). Tiller number was approximately 15 across WT and *pin2* with an average of 5 leaves per tiller (Figure S2). Similarly, leaf size was consistent across mutant and wild type in both rhizobox and seedling assays (Figure S1).

A maturity study was conducted to explore canopy traits at maturity, where no significant changes to above-ground canopy biomass, number of fertile tillers at maturity and days to flowering were observed. Following spatial analysis, no genotypic effects were associated with these traits between WT and *pin2* (Figure S3).

## DISCUSSION

### *PIN2* is essential for root gravitropism and root angle

The spatial and temporal distribution of roots via branching, elongation and curving all affect the ability of the crop to extract water and nutrients from the soil (Manschadi et al. 2006; Rich and Watt 2013). A substantial body of research suggests that optimised RSA can improve water acquisition with minimal metabolic cost, potentially improving crop yields under both favourable and adverse conditions (Uga 2021). Specifically, variation in gravitropism can affect root growth angle, altering root anchoring and exploration of different soil layers to acquire water and nutrients, as shown for *ENHANCED GRAVITROPISM 2* (*EG2*) in barley and wheat (Kirschner et al. 2021). However, our knowledge of all the genes involved in orchestrating root growth and gravitropism is limited, particularly in cereals. Here we report that Hv-*PIN2* plays an important role in gravitropism and root angle in barley. Plants with a loss of *PIN2* display root-specific agravitropism, that increased root angle resulting in wider and shallow rooting, without impacting the above-ground traits. These findings demonstrate the potential for *PIN2* as a new target to improve climate resilience, while maintaining the agronomic traits of the canopy that have already been optimised through breeding for different crop production regions.

Deep root systems can contribute to drought adaptation by improving access to stored soil moisture in deep soil layers, thereby providing an advantage during terminal or post-flowering drought stress. The root system of the crop can exhibit morphological, structural and physiological responses to changes in the growing environment, referred to as root developmental plasticity. This is underpinned by changes in root branching, elongation and curving which broadly define the extent to which the root system can explore the soil volume. The ‘curving’ component is associated with angle of seminal roots, and the seedling proxy trait of mature root angle (Kang et al. 2024).

Despite the more complex genetic and physiological architecture of barley, our analysis of tropic responses through *pin2* knockouts revealed homologous function to that of the model species Arabidopsis (Müller et al. 1998). Typically, gravitropism is a guiding force whereby the radicle grows directly downwards, followed by a slight weakening of the response to initiate growth at non-vertical angles in order to facilitate roots spreading for soil exploration (Sachs and Botanik 1882). This is termed the gravitropic setpoint angle and is considered a balance between two antagonistic growth signals, gravitropism and a currently unidentified anti-gravitropic offset (Digby and Firn 1995). In *pin2* knockouts, the roots appeared ‘aimless’, and instead of exhibiting a consistent growth angle in response to gravity as typically seen in seedling assays, the roots continued to grow from their point of initiation out of the soil, against the gravity vector. In addition, phenotyping seedlings for seminal root angle (an established proxy for mature root angle) showed significant increases in seminal root angle compared with WT (Rambla et al. 2022; Richard et al. 2015). These observations suggest the loss of gravistimulation leads to a loss of the primary guiding force for root angling, one of the earliest drivers of root distribution and architecture formation (Roychoudhry et al. 2013; Digby and Firn 1995; Nakamoto and Oyanagi 1996).

There is a well-established link between root distribution in the soil, water access and water use (Robertson et al. 1993; Li et al. 2022). Further evidence on the role of *PIN2* and gravitropism in directing RSA formation in barley comes from functional analysis. The increased root angle resulting from inhibited apical auxin transport caused a wider and shallower root system in *pin2*, resulting in larger root biomass in upper layers, while trending towards a reduction in roots at depth, resembling a topsoil foraging ideotype that is beneficial for capturing phosphorous and intermittent rainfall (Lynch 2013, 2019).

The identification of auxin-related mutants in model species like Arabidopsis has played a key role in better understanding how auxin orchestrates plant morphogenesis. For example, loss of At-*PIN2* causes root gravitropism defects due to the mutant’s inability to transport auxin from the roots towards the shoots and, due to *PIN2*s high levels of conservation in cereals, a similar function is predicted in cereals that exhibit more complex root architecture (Lu et al. 2015).

### *PIN2* mutants cause root system-specific changes

The quiescent centre is a group of up to 1000 cells located in the root tip (apical meristem) of vascular plants where cell division occurs (CLOWES 1958). In Arabidopsis, At-PIN1 acts in shoot to root auxin transport, moving auxin towards the quiescent centre of the root meristem (Blilou et al. 2005), and At-PIN2 functions basally of the quiescent centre and meristem to inhibit cell elongation in the root tip. Based on the existing literature, we hypothesized that a build-up of auxin in the lateral root cap post-quiescent centre in *pin2* mutants is the cause of the root-specific changes to cell expansion, while *pin1* mutants, as they act pre-quiescent centre, affect cell division and distal patterning (Forestan and Varotto 2010; Kirschner et al. 2018). Overall, this downstream effect on root elongation allows for more subtle changes to root architecture without significantly affecting organism-wide patterning and architecture. For example, the overexpression of Zm-*PIN1a* showed an elevated concentrations of auxin in the roots, increasing lateral root formation and seminal root length, improving root weight and shortening plant height (Forestan and Varotto 2010). Typically, auxin in the WT is transported towards the roots from their site of synthesis in the leaf primordia (Figure S4). Once transported to the root tip by PIN1, auxin accumulates at the root meristem, inhibiting elongation (Swarup et al. 2005). Subsequently, PIN2 transports auxin along the apical side of the roots, shootward (Blilou et al. 2005). This PIN2 transport along the base of the root inhibits the roots elongation and causes the roots to curve downwards, also known as the gravitropic response. Auxin is then transported back to the root tip for the process to repeat. In the *pin2* knockout lines, we hypothesise the root tips are accumulating auxin due to an inability to flush auxin away without the apical shootward auxin transport typically facilitated by PIN2 (Figure S4). Thus, there is limited gravistimulation leading to increases in root angle and causing a visible agravitropic phenotype.

Based on our findings, Hv-*PIN2* has been shown to be a root-specific auxin efflux transporter required for gravitropic responses in the roots. Loss of function produced ‘aimless’ rooting with agravitropism, leading to increased root angle and a shift towards a top-soil foraging RSA with a potential increase in root biomass (Figure 4). Interestingly, under the well-watered conditions used in this study, the *pin2* barley mutant did not show any difference in above-ground biomass. This is consistent with the overexpression of the Os-*PIN2* homologue in rice, where no significant changes to canopy branching (tiller number), leaves per tiller, area per leaf, number of fertile tillers or days to flower were observed (Chen et al. 2012). The lack of changes to canopy architecture or flowering time in cereal crops suggests this root-specific target could be directly used in the future to breed cultivars with shallow rooting systems. Summarizing, Hv-*PIN2* has been characterized as the functional homologue of At-*PIN2*, playing an essential role in root gravitropism and providing new insight into root and shoot auxin transport and its roles in development and gravitropism.

### Root:shoot ratio of *pin2* mutants is insensitive to planting density

Genetic and management solutions are required to develop resilient crops in highly variable environments. Many different management systems are possible to combat drought such as row spacing, population, irrigation, fertilization and cropping systems. Many different genotypic solutions are also possible, including utilizing the PIN gene family to modify traits affecting water supply and demand. Understanding the interaction of genotypes with different management strategies for specific target environments is critical to improving yield stability. While *PIN2* clearly plays an important role in orchestrating root curving through gravitropism, it also has novel effects on allometry under different planting densities in glasshouse conditions. Plants with narrow angle and more vertical roots tend to have larger proportions of their roots at depth during later development which could increase access to water in deep soils. Conversely, genotypes with wider root angle and more horizontal rooting are better able to explore the soil in the inter-row space which could increase access to water in skip-row systems (Hammer et al. 2009).

In fertile environments when water is not limiting, competition among plants in a crop is mainly for light. However, in nutrient-poor or water-scarce conditions, competition is mainly for the limiting factor of either nutrients or water, respectively. The effects of competition among root systems are largely determined by the levels of water and nutrient availability, particularly inorganic forms of nutrients such as nitrate and ammonium (Tilman 1985). The usual row spacing of barley is typically 15-20 cm between rows to maximise the potential of individual plants. In our rhizobox experiments with non-limiting water and nutrients, planting density (number of plants per rhizobox) was positively correlated with RSRatio in the WT lines. As plants per box increased from 0.5 (30 cm row spacing) to 1 (15 cm row spacing) to 3 (5 cm row spacing), the RSRatio increased from 0.3 to 0.5 to 0.7, respectively. This indicates an increased allocation of resources to the roots as planting density increased, likely a cost in response to increased above-ground competition. Whereas in contrast, the *PIN2* mutant lines maintained a RSRatio of about 0.4 regardless of planting density, highlighting a trending insensitivity of *pin2* to competition i.e. a reduced or loss of sensitivity to rooting density on root growth. Hence, at higher planting densities there was a significant change in root:shoot biomass ratio, with no significant change to canopy biomass and a reduced root system, consistent with the *pin2* response in Arabidopsis and likely a reflection of the reduction in root elongation (Figure S1, Muller., 1998). Field trials will be necessary to fully explore these implications on crop performance across a range of conditions.

In the glasshouse environment used in the rhizobox experiments, nutrient availability was not a limiting factor. As such nutrient uptake or the diffusion rate of the ions in the soil may have been limiting factors. While the WT reflexively altered its biomass allocation depending on planting density, increasing below-ground biomass as planting density increased, *pin2* maintained a consistent RSRatio at all tested densities (Figure 4D). As such, it is hypothesized *PIN2* plays an important role in external sensing alongside gravitropism, where the absence of this PIN2 transporter leads to agravitropic roots that project outwards and appear insensitive to environmental stimuli such as intraspecific competition. In field grown barley when nutrients are not limiting, less biomass is required underground and capacity for nutrient uptake can be enhanced by concentrating root proliferation in the topsoil (GRIME and HODGSON 1987; Hecht et al. 2016). It is possible the increased root proliferation in upper soil layers by *pin2* provides improved resource availability limiting the need for further resource investment into soil exploration. Therefore, modulating the expression of *PIN2* provides a unique target for improving RSA, which may lead to a range of root angles and influence drought tolerance without any negative impacts on yield or obvious above-ground trade-offs (Joshi et al. 2016). Currently, the complete loss-of-function *pin2* mutants display a topsoil foraging root system ideotype that may provide benefits for uptake of immobile nutrients in some agricultural settings. However, field studies in different environments are needed to explore the effects of *PIN2* on yield and its influence under changing planting densities (Liu and Godwin 2012).

Understanding the allometric relationship between canopy and root architecture has important agricultural implications. Here we demonstrate conserved function of the auxin efflux carrier Hv-*PIN2* in directing root angle through the gravitropic response in barley. Alongside altered root distribution that produced a shallower and more expansive RSA, *pin2* mutants did not display the typical root-shoot allometric relationship. While further investigation is required regarding the exact mechanism regulating root angle, *PIN2* shows clear links to agronomic traits such as root angle and root distribution that may offer an advantage under moisture-limited environments. The novel root phenotypes identified in this study offer useful insight into the potential for genome engineering of root traits for future barley improvement programs. Overall, the investigation of key developmental genes (initially discovered in model species) through gene editing shows potential to support the development of new cultivars with improved nutrient-use efficiency and climate resilience.

## AUTHOR CONTRIBUTIONS

K.M. conceived the study. K.M. design, synthesis, and transformation of gene edits. Z.A. analysed the transgenics, including phenotyping experiments and data analyses. H.R guided assisted experimental design and analysis of data. All authors were involved in interpretation of datasets and results. R.D assisted in phenotyping, particularly the double width rhizobox platform. Z.A. wrote the manuscript, and all authors were involved in editing and approved the manuscript.

## ACKNOWLEDGMENTS

We are grateful to the Crisp group for stimulating discussions during the work. The authors are also grateful to Dr Millicent Smith and Dr Owen Powell at The University of Queensland for coordinating review and guiding constructing of the early stages of the manuscript. This research was supported by ARC Discovery Project ‘Cereal blueprints for a water-limited world’ (DP190102185) and ARC Linkage project ‘Digging deeper to improve yield stability’ (LP200200927). Z.A. received scholarships from the Australian Government Research Training Program and InterGrain. L.H. was supported through an ARC Future Fellowship (FT220100350).

## SUPPORTING INFORMATION

**Figure S1.** Illustration of *PIN2* genome edited knock out lines with impact on protein structure and transmembrane domains compared to wild type.

**Figure S2.** Canopy and root trait results from narrow rhizobox experiments conducted using different planting density.

**Figure S3.** Seedling and maturity trials reveal no significant difference to canopy and phenology in *pin2.*

**Figure S4.** Summary of hypothesis: disrupted auxin transport in *pin2* genome edited barley inhibits gravitropic response and alters root development.

**Table S1.** Fixed and random effects table for the linear mixed model (LMM) fitted to the trait of interest.

**Table S2.** Synteny of *PIN* genes across cereals.

---

**Supporting figure 1.** Illustration of *PIN2* genome edited knock out lines with impact on protein structure and transmembrane domains compared to wild type.

Predicted transmembrane domains (TMDs) of wild type and transgenic lines using membrane protein topology prediction method TMHMM, based on a hidden Markov model. Peaks represent TMDs, with the x axis corresponding to the position in the protein sequence, and the y axis representing the probability of the model as a representative scale for confidence in prediction. Blue lines represent intracellular protein domains, red represents TMDs and pink is extracellular regions. LBS = Ligand Binding Site, G1 and G2 = Guide sites one and two for CRISPR/Cas9 genomic editing, TMD = Transmembrane Domain. Further description of edits outlined in Figure 3.

**Supporting figure 2.** Canopy and root trait results from narrow rhizobox experiments conducted using different planting density.

**A)** Comparative data for 3 and 1 plant per box rhizobox (30 cm x 5 cm x 80 cm) trials. Comparisons for total biomass, above-ground biomass, below-ground biomass, the root:shoot biomass ratio (RSRatio)

**B)** Representative image of single plant per box, whole rhizobox (30 cm x 5 cm x 80 cm) comparison between control WT and *pin2-1* knockout line. *pin2-1* displayed no visual reduction in canopy size compared with WT despite changes in the RSA.

**C)** Root:Shoot biomass ratio for each section of the rhizobox where upper, middle, lower equal S1, S2 and S3 respectively. N=8 for each genotype in each trial. Data analysis was performed using ASReml and ASReml Plus for spatial analysis, calculation and comparison of BLUPs. Standard parametric t-test was used to calculate significance between planting densities. ns = P>0.1, * = P<0.05, • = P<0.1

**Supporting figure 3.** Seedling and maturity trials reveal no significant difference to canopy and phenology in *pin2*.

**A)** Seedling assay (N=14) comparisons between wild type (WT) and *pin2* edited lines (*pin2-1* and *pin2-2*) of total root:shoot biomass ratios (RSRatio) and average leaf area (3 technical replicates) for the number of leaves on the main stem.

**B)** Phenology and canopy measurements from a maturity study (N=8). Seedling assay and maturity study identified no significant change to above ground canopy architecture despite significant decrease in below ground biomass. No genotypic correlation to differences in canopy biomass (g), number of fertile tillers and days to flower traits at maturity. Data analysis was performed using ASReml and ASReml Plus for spatial analysis, calculation and comparison of BLUPs. Error bars display standard error of spatially adjusted mean. ns = P>0.1, * = P<0.05, ** = P<0.01

**Supporting figure 4.** Summary of hypothesis: disrupted auxin transport in *pin2* genome edited barley inhibits gravitropic response and alters root development.

Wild type (*PIN2*) root tip illustrating auxin transport and accumulation at the meristem in orange. Overall generalised wild type root architecture displayed on the left. Auxin is transported towards the roots from their site of synthesis in the leaf primordia. Once transported to the root tip by PIN1 they accumulate at the root meristem, inhibiting elongation. PIN2 transports auxin along the apical side of the roots, shootward. This PIN2 transport along the base of the root inhibits the roots elongation and causes the roots to curve downwards, also known as the gravitropic response. Auxin is then transported back to the root tip for the process to repeat. The *pin2* lines accumulate auxin at the root tip due to inability to flush auxin away. Without the apical shootward auxin transport facilitated by PIN2, there is limited gravistimulation, increasing root angle and causing a visible agravitropic phenotype. Design was adapted from Dhonukshe, 2012 and auxin distribution in *PIN2* KO was adapted from Thomas, 2023.

**Supporting table 1.** Fixed and random effects table for the linear mixed model (LMM) fitted to the trait of interest.

Each trait fit a custom LMM measuring across glasshouse root and shoot experiments. Broad sense heritability for each trait was calculated to measure the proportion of phenotypic variance attributed to genetic effects.

**Supporting table 2.** Synteny of *PIN* genes across cereals.

Homology was determined using Phytozome comparing protein sequence homology and genomic sequence (Goodstein et al. 2012). Alternate naming conventions are included where the common name differs.

