## Supplementary material for "The *PIN2* ortholog in barley modifies root gravitropism and architecture without impacting the shoot": Table S1

| *Experiment* | *Trait* | *Fixed Effects* | *Random Effects* | *Broad sense heritability* |
|---|---|---|---|---|
| *Maturity Study* | Tiller number | lin(Column) + lin(Row) | Genotype + Rep | 7.97E-04 |
| | Fertile Tiller number | lin(Row) | Genotype + Rep + Row | 2.96E-04 |
| | Days to Flowering | | Genotype + Rep | 1.49E-05 |
| | Canopy Biomass | lin(Column) + lin(Row) | Genotype + Rep | 1.12E-05 |
| *Rhizobox* | Below-ground Biomass | | Genotype + Block + Row | 3.87E-01 |
| | Above-ground Biomass | lin(Column) | Genotype + Block + Row | 4.84E-01 |
| | Total Biomass | lin(Column) | Genotype + Block | 1.69E-01 |
| *Seedling Assay* | Shoot Biomass | lin(Column) + lin(Row) | Genotype + Block + Bench | 9.20E-01 |
| | Root Biomass | lin(Column) + lin(Row) | Genotype + Block | 8.25E-01 |

**Supporting table 1.** Fixed and random effects table for the linear mixed model (LMM) fitted to the trait of interest.
