## Supplementary material for "The *PIN2* ortholog in barley modifies root gravitropism and architecture without impacting the shoot": Table S2

| Arabidopsis | AtCode | Rice | Os_LOCcode | OsCode | Sorghum | SobicCode | Maize | GRMZcode | ZmCode(v4) | Barley | HORVUcode |
|---|---|---|---|---|---|---|---|---|---|---|---|
| AtPIN1 | At1g73590 | PIN1A | LOC_Os06g12610 | Os06g0232300 | SbPIN10 | Sobic.010G095100 | ZmPIN1a | GRMZM2G098643 | Zm00001d044812 | PIN1A | HORVU7Hr1G038700 |
| | | PIN1B | LOC_Os02g50960 | Os02g0743400 | SbPIN6 | Sobic.004G240900 | ZmPIN1b | GRMZM2G074267 | Zm00001d018024 | PIN1B | HORVU6Hr1G076110 |
| | | PIN1C | LOC_Os11g04190 | Os11g0137000 | SbPIN7 | Sobic.005G025900 | ZmPIN1c | GRMZM2G149184 | Zm00001d052269 | PIN1C | HORVU4Hr1G026680 |
| | | PIN1D | LOC_Os12g04000 | Os12g0133800 | | | ZmPIN1d (SoPIN1) | GRMZM2G171702 | Zm00001d052442 | PIN1D | |
| AtPIN2 | At5g57090 | PIN2 (LRA1) | LOC_Os06g44970 | Os06g0660200 | SbPIN11 | Sobic.010G216700 | ZmPIN2 | GRMZM2G074267 | Zm00001d018024 | PIN2 (LRA1) | HORVU7Hr1G110470 |
| AtPIN3 | At1g70040 | n/a | n/a | n/a | n/a | n/a | n/a | n/a | n/a | n/a | n/a |
| AtPIN4 | At2g01420 | n/a | n/a | n/a | n/a | n/a | n/a | n/a | n/a | n/a | n/a |
| AtPIN5 | At5g16530 | PIN5A (PIN6) | LOC_Os01g69070 | Os01g0919800 | SbPIN5 | Sobic.003G403900 | ZmPIN5a | GRMZM2G025742 | Zm00001d042345 | PIN5A (PIN6) | HORVU3Hr1G094000 |
| | | PIN5B | LOC_Os08g41720 | Os08g0529000 | SbPIN8 | Sobic.007G195201 | ZmPIN5b | GRMZM2G148648 | Zm00001d031594 | PIN5B | HORVU5Hr1G076210 |
| | | PIN5C | LOC_Os09g32770 | Os09g0505400 | SbPIN1 | Sobic.002G257001 | ZmPIN5c | GRMZM2G040911 | Zm00001d006082 | PIN5C | HORVU5Hr1G076060 |
| AtPIN6 | At1g77110 | n/a | n/a | n/a | n/a | n/a | n/a | n/a | n/a | n/a | n/a |
| AtPIN7 | At1g23080 | n/a | n/a | n/a | n/a | n/a | n/a | n/a | n/a | n/a | n/a |
| AtPIN8 | At5g15100 | PIN8 | LOC_Os01g51780 | Os01g0715600 | SbPIN3 | Sobic.003G276700 | ZmPIN8 | GRMZM5G839411 | Zm00001d043660 | PIN8 | HORVU3Hr1G067670 |
| monocot sp | n/a | PIN9 | LOC_Os01g58860 | Os01g0802700 | SbPIN4 | Sobic.003G327500 | ZmPIN9 | GRMZM5G859099 | Zm00001d043179 | PIN9 | HORVU3Hr1G078620 |
| monocot sp | n/a | PIN10A | LOC_Os01g45550 | Os01g0643300 | SbPIN2 | Sobic.003G235800 | ZmPIN10a | GRMZM2G126260 | Zm00001d044083 | PIN10A | HORVU3Hr1G057630 |
| | | | | | | | | | | PIN10B-1 | HORVU1Hr1G091030 |
| monocot sp | n/a | PIN10B | LOC_Os05g50140 | Os05g0576900 | SbPIN9 | Sobic.010G051000 | ZmPIN10b | GRMZM2G160496 | Zm00001d045219 | PIN10B-2 | HORVU1Hr1G072970 |
| | | | | | | | | | | PIN10B-3 | HORVU3Hr1G111420 |

**Supporting table 2.** Synteny of *PIN* genes across cereals.
