## Supplementary material for "The *PIN2* ortholog in barley modifies root gravitropism and architecture without impacting the shoot": Figure S1

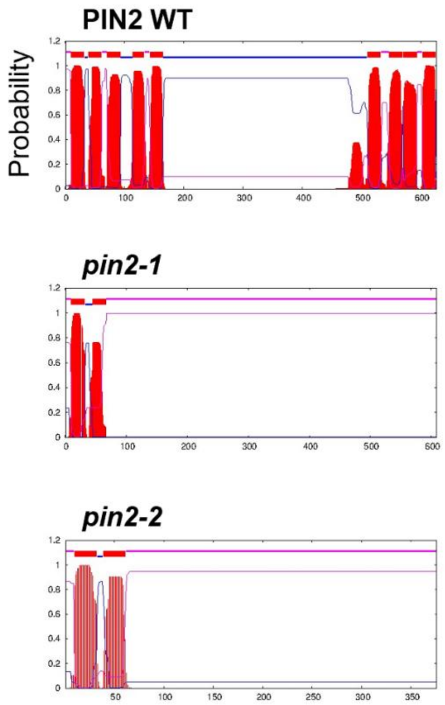


**Supporting figure 1.** Illustration of *PIN2* genome edited knock out lines with impact on protein structure and transmembrane domains compared to wild type.
