## Supplementary material for "The *PIN2* ortholog in barley modifies root gravitropism and architecture without impacting the shoot": Figure S2

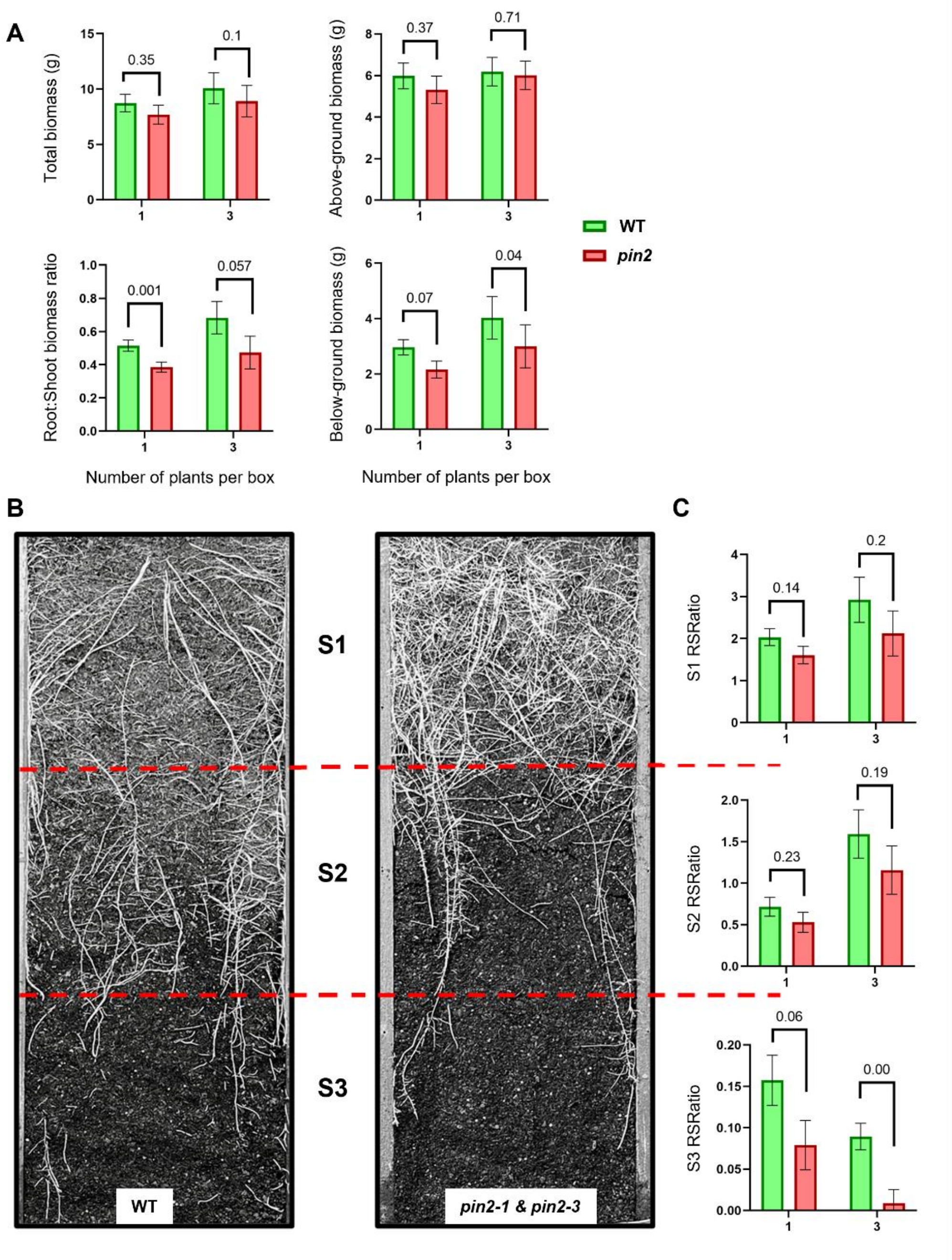


**Supporting figure 2.** Canopy and root trait results from narrow rhizobox experiments conducted using different planting density.
