## Supplementary material for "The *PIN2* ortholog in barley modifies root gravitropism and architecture without impacting the shoot": Figure S3

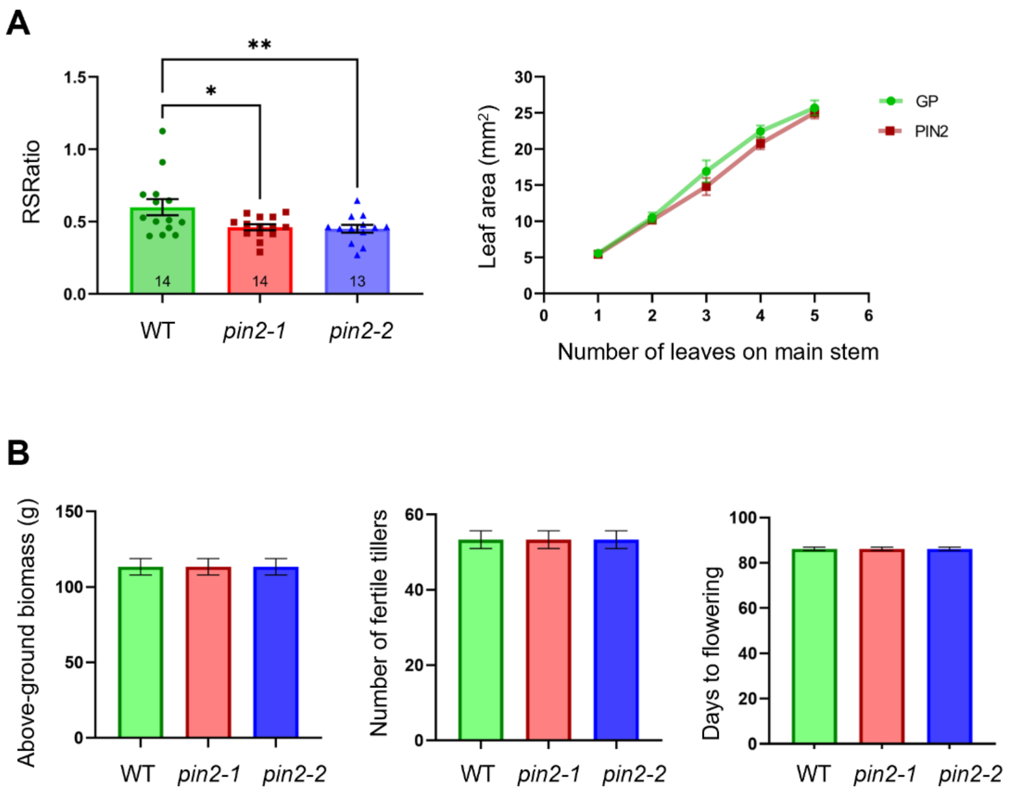


**Supporting figure 3.** Seedling and maturity trials reveal no significant difference to canopy and phenology in *pin2*.
