## Supplementary material for "The *PIN2* ortholog in barley modifies root gravitropism and architecture without impacting the shoot": Figure S4

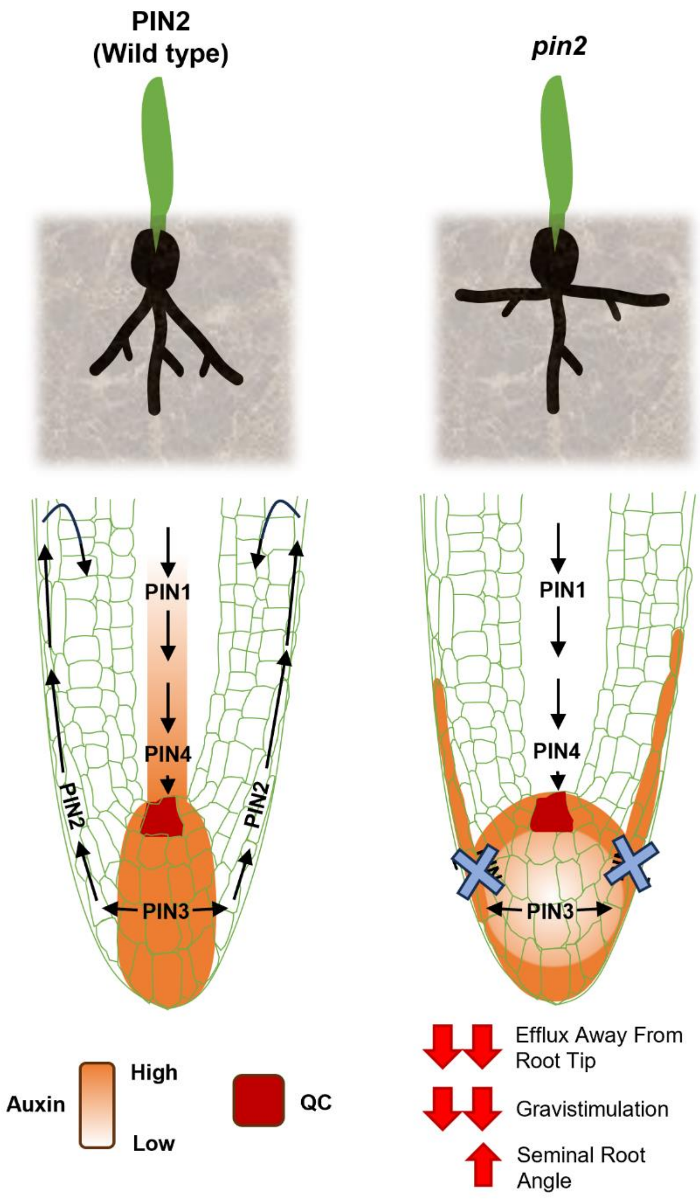


**Supporting figure 4.** Summary of hypothesis: disrupted auxin transport in *pin2* genome edited barley inhibits gravitropic response and alters root development.
